# Closed-loop and automatic tuning of pulse amplitude and width in EMG-guided controllable transcranial magnetic stimulation (cTMS)^⋆^

**DOI:** 10.1101/2022.05.08.491097

**Authors:** S. M. Mahdi Alavi, Fidel Vila-Rodriguez, Adam Mahdi, Stefan M. Goetz

## Abstract

This paper proposes a tool for automatic and optimal tuning of pulse amplitude and width for sequential parameter estimation (SPE) of the membrane time constant and input–output curve in closed-loop electromyography-guided (EMG-guided) controllable transcranial magnetic stimulation (cTMS). A normalized depolarization factor is defined which separates the optimization of the pulse amplitude and width. Then, the pulse amplitude is chosen by the maximization of the Fisher information matrix (FIM), while the pulse width is chosen by the maximization of the normalized depolarization factor. The simulation results confirm satisfactory estimation. The results show that the normalized depolarization factor maximization can identify the critical pulse width, which is an important parameter in the identifiability analysis, without any prior neurophysiological or anatomical knowledge of the neural membrane.

## 1. Introduction

Transcranial magnetic stimulation (TMS) is a therapeutic intervention used in psychiatric and neurological disorders as well as a tool for probing brain function in cognitive neuroscience and neurophysiology [1, 2, 3, 4]. In parallel to the rapid spread of TMS usage, novel technological refinements and developments are emerging, such as TMS devices with adjustable pulse shapes, among them controllable TMS (cTMS), and closed-loop TMS [5, 6, 7, 8, 9].

The cTMS device allows adjustment of the pulse width [10, 11]. It enables optimization, selectively targeting distinct neuronal populations, and quantification of the strength–duration and input–output (IO) behaviors and curves [12, 13, 14, 15, 16, 17]. Different pulse shapes, including the pulse duration can influence the balance between concurrently activated neural elements. For example, brief pulses, i.e., pulses with a short duration or width, may result in preferential motor than sensory stimulation in peripheral nerves [18]. Studies demonstrated that the ratio of cortical motor to scalp sensory threshold is lower for brief pulses compared to longer stimuli [19]. It was found that short duration pulses inducing dominant anterior–posterior (AP) currents enable the selective recruitment of the longest latency motor evoked potentials (MEPs) compared to posterior–anterior (PA) and long duration AP currents [12, 13]. In contrast, the longer pulse width leads to higher local mean field power (LMFP) according to the TMS–electroencephalography (TMS–EEG) study in [20]. It was shown that the longer pulse widths result in lower motor thresholds and steeper IO curves [22]. The pulse width may also impact tolerability as research has shown that wider TMS pulses are more tolerable and are associated with less discomfort [23].

Despite ignoring the strong nonlinearity, the so-called time constant, which approximates neurons as a first-order low-pass filter with a specific time constant *τ*_*m*_, is widely used metric to characterize the activation dynamics of neurons. Although this approach may be inappropriate to simulate neurons and predict pulse shape effects, it is still very sensitive for detecting changes. Despite the limitations, the time constant differs between various targets and can identify the activated neuronal target [12, 24]. Furthermore, it has been suggested and tested for diagnosis [25, 26, 27, 28].

The detection of the time constant as the key parameter of the simplified linear approximation of the activated neural target needs a detectable response, which is more readily available for stimulation in the visual cortex and the (primary) motor cortex [29, 30]. Stimulation in the motor cortex directly or indirectly activates corticospinal fibers that send the signal to the periphery, where it can be detected as muscle contractions [31]. Although these motor responses can in principle be detected visually, electromyography (EMG) provides quantitative measures and is preferred as it allows substantially higher accuracy and sensitivity for response detection [32, 33, 34]. The measurement as well as analysis of an EMG signal to extract response information online, during a procedure and adjustment of the procedure based on that extracted information started the field of so-called closed-loop TMS.

Closed-loop TMS refers to the automatic and real-time adjustment of TMS parameters based on measurements, e.g., to maximize the desired plastic effects by using the brain/neural data in a feedback system or to speed up the detection of a biomarker [35, 36]. Closed-loop TMS is an area of active research using both closed-loop EEG-guided TMS [36, 37, 38, 39, 40] and closed-loop EMG-guided TMS [46, 47, 48].

In [36], an algorithm was developed to determine the electric-field orientation that maximizes the amplitude of the TMS–EEG response. References [37] [38] [39], and [40] address timing of TMS pulses by using real-time EEG to target phases of ongoing oscillations that are associated with high excitability states of the neuronal networks.

Applications of the closed-loop EMG-guided TMS include automatic and optimal estimation and characterization of the neural membrane and input–output (IO) behaviors. IO curves show large nongaussian variability so that their analysis typically requires appropriate mathematical treatment against bias, while closed-loop procedures enable faster estimation [41, 42, 43, 44, 45]. Optimal estimation of the IO and slope curves by using Fisher-information-based sequential parameter estimation (FIM SPE) methods has been addressed in [46] and [47]. In the proposed FIM SPE methods, the prior knowledge of the neural membrane and input–output (IO) behaviors are analyzed in a closed-loop system for the selection of the next pulses, which result in more accurate estimations, with fewer numbers of TMS pulses, reducing the duration and discomfort of the procedure, compared to uniform sampling methods. It is re-called that the uniform-sampling-based estimation is an open-loop system, where the TMS stimuli are uniformly distributed between the minimum and maximum values [49, 50, 51]. There exists a main drawback that there is no specific or systematic approach to optimally choose the number and order of the TMS stimuli in the uniform-sampling-based estimation.

In [48], an integrated model of the neural membrane time constant and IO curve was developed by analytical computation of the depolarization factor in cTMS. A FIM SPE method was presented for the estimation of the neural membrane time constant and IO curve at fixed pulse widths shorter than the critical pulse width where the peak of the membrane response occurs at the end of the pulse. The peak of the membrane response occurs before the end of the pulse for all pulse widths longer than the critical pulse width. However, the pulse widths are picked manually and rather ad hoc, e.g., randomly, and only, the pulse amplitude is adjusted automatically using the FIM SPE [48]. Therefore, the proposed FIM SPE method in [48] is not optimal in terms of the pulse-width selection.

This paper focuses on automatic and optimal tuning of the pulse width for optimal estimation of the neural membrane time constant and IO curve. A computational algorithm is developed which consists of two tuning stages. The pulse width is adjusted by maximizing the normalized depolarization factor, and the pulse amplitude is optimally chosen by using the FIM SPE method as per [46]. In [48], the pulse width should be adjusted using the empirical critical pulse width data. This is not needed for the proposed method in this paper. The results also show that the depolarization factor maximization computes the critical pulse width with a satisfactory level of accuracy.

The contributions of this paper are then highlighted as follows:

– Formalism for automatic and optimal tuning of both the pulse amplitude and width in cTMS for optimal estimation of the neural membrane time constant and IO curve.
– Determination of the critical pulse width by depolarization factor maximization.

## 2. The neural system model

The proposed cTMS pulse tuning method is based on the following integrated model of the neural time constant and IO curve [48]

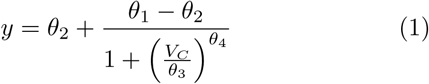

where

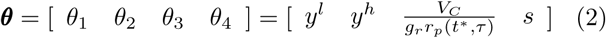

is the parameter vector. *V*_*C*_ denotes the normalized pulse amplitude adjustable between *V*_*C*_(min) = 0 and *V*_*C*_(max) = 1. *y i*s a characteristic of MEP such as its peak-to-peak value, area under the curve (AUC), root mean square or latency [51]. This paper chooses the MEP’s peak-to-peak value as *y*. The parameters *y*^*l*^ and *y*^*h*^ denote the lower and upper plateaus of the IO curve. *τ d*enotes the membrane time constant.

The parameter *r*_*p*_(*t*_*p*_, *τ)* denotes the depolarization factor which is defined as

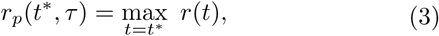

where *r*(*t*) is the neural response to the cTMS pulse, and *t*^*∗*^ is the peak time.

The linearized neural response estimate as typically used for the time constant is obtained by the convolution of the cTMS pulse and the neural membrane dynamics. For the cTMS pulse class given in [10] and the first-order neural membrane dynamics

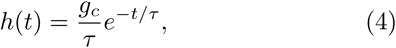

the linearized neural response is given by

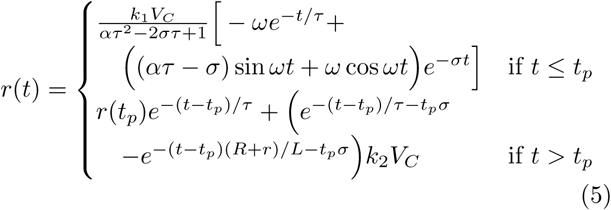

where *t*_*p*_ is the pulse width [48]. It is assumed to be adjustable between *t*_*p*_(min) = 10 µs and *t*_*p*_(max) = 200 µs. *r*(*t*_*p*_) is the neural membrane response at *t* = *t*_*p*_, and

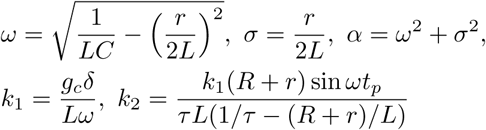

*L* is the stimulating coil, *R* is the energy dissipation resistor, *r* is the resistor which represents the combined resistance of the capacitor, inductor and semi-conductor switch. The following parameter values are used in this paper [10]: *L* = 16 µH, *C* = 716 µF, *R* = 0.1 Ω, *r* = 20 mΩ, *δ* = 3.2 × 10^−6^ (V/m)(A/s).

The depolarization factor is computed as follows:

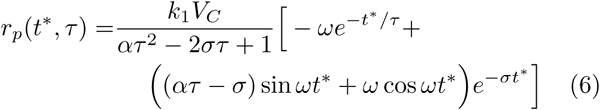

The gain *g*_*c*_ characterizes the coupling between the TMS coil and the directly stimulated neurons and hence depends on factors such as the coil-to-cortex distance, head size, and neural population type and orientation relative to the induced electric field. The *r*_*p*_ value is small, thus the gain *g*_*r*_ is utilized to scale it up. The mid-point and slope of the IO curve are given by *r*_*p*_ and *s*.

To separate the tuning of the pulse amplitude and width, the following normalized depolarization factor is defined:

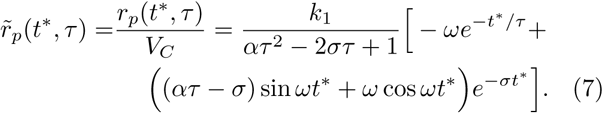

Fig. 1 illustrates a sample cTMS pulse, neural response and depolarization factor.

**Figure 1:**
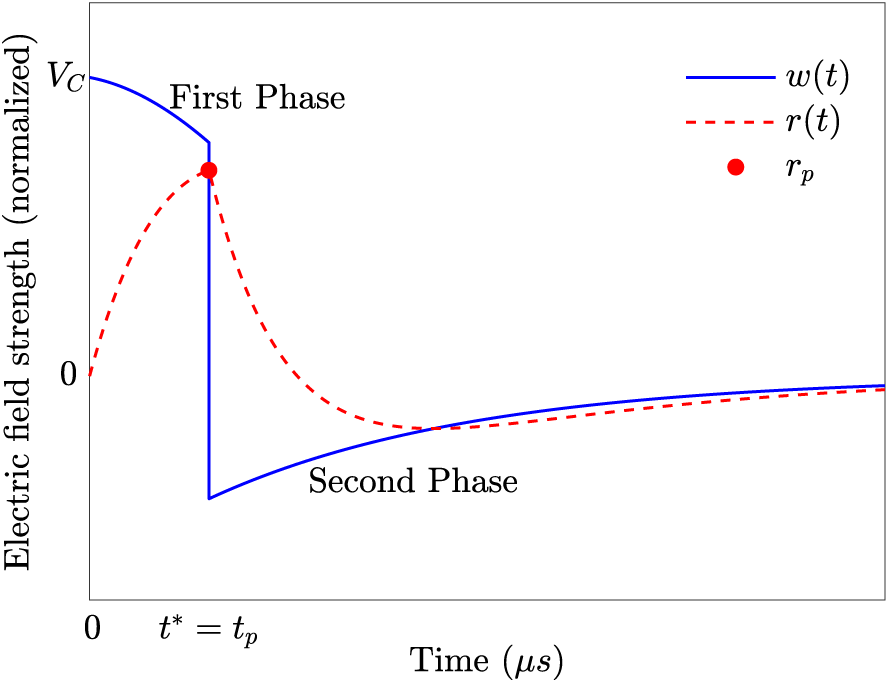
Sample cTMS pulse, neural response and depolarization factor.

The depolarization factor *r*_*p*_ occurs either at the end of the pulse (*t*^∗^ = *t*_*p*_) or before the end of the pulse (*t*^∗^ < *t*_*p*_). The critical pulse width 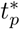 is defined as the pulse width for which *r*_*p*_ occurs at *t*^∗^ = *t*_*p*_ for all pulse widths shorter than that, and *r*_*p*_ occurs at *t*^∗^ < *t*_*p*_ for all pulse widths longer than that. The values of *t*^∗^ and 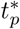 depend on the the pulse parameters and *τ*.

Fig. 2 shows the critical pulse width versus the time constant. By fitting to the analytical data, the following relationship is obtained between 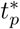 and *τ* :

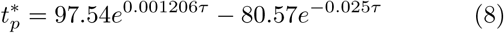

**Figure 2:**
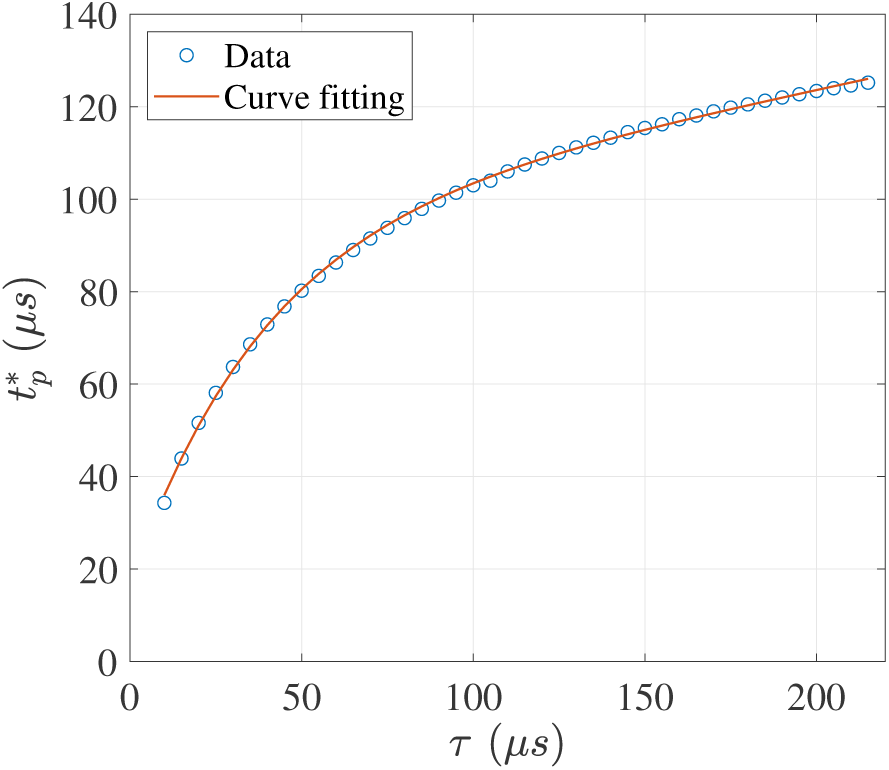
The critical pulse width versus the time constant in cTMS.

It was shown that the integrated model (1) is identifiable for 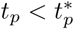 [48].

## 3. Problem Statement

The problem is to design an algorithm for automatic and optimal tuning of the cTMS pulse amplitude and width to estimate the integrated model parameters, i.e., neural membrane time constant *τ* and the IO curve parameters *θ*_*i*_, *i* = 1, …, 4 in a closed-loop system.

In [48], a pulse width *t*_*p*_ shorter than the critical pulse width is chosen manually by using Fig. 2, which requires a prior knowledge of the neural time constant. However, the exact time constant of the neural membrane is unknown in practice. The proposed technique in this paper does not require prior information of the time constant or critical pulse width.

## 4. Proposed algorithm for automatic and optimal tuning of cTMS pulse parameters

Fig. 3 shows the functional block diagram of the proposed algorithm, which is based on the sequential parameter estimation (SPE) method.

**Figure 3:**
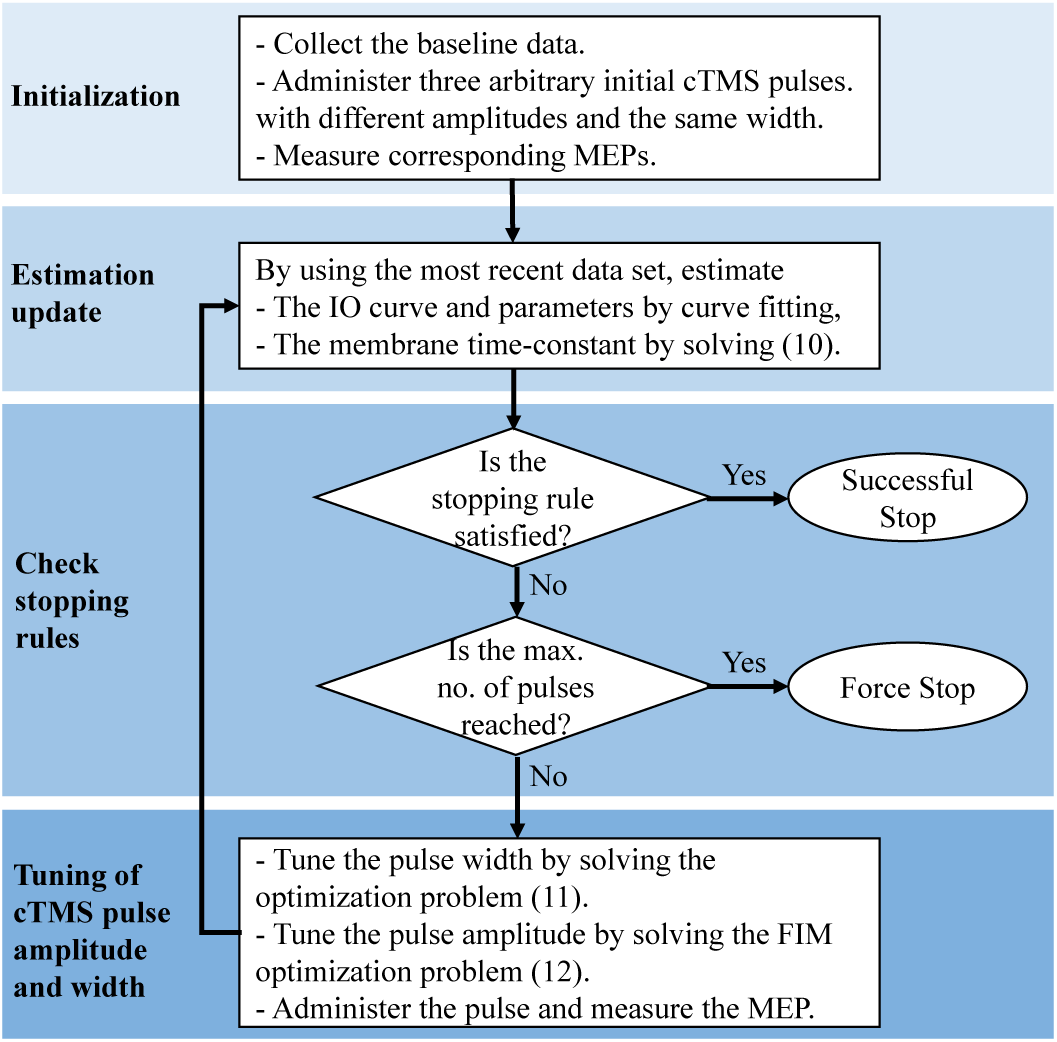
Functional block diagram of automatic and optimal tuning of pulse amplitude and width in closed-loop EMG-guided cTMS.

The proposed SPE algorithm consists of four main parts: initialization, selection of estimation update, estimation update and checking the stopping rule.

In the initialization stage, the baseline data is firstly collected. Then, three initial cTMS pulses are administered with the magnitudes 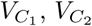, and 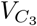 chosen arbitrarily between *V*_*C*_(min) and *V*_*C*_(max). The width of the initial pulses should be fixed, i.e., 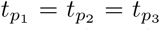. According to the identifiability analysis in [48], the pulse width should be shorter than or equal to the critical pulse width 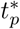, i.e., 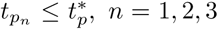. If 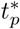 is unknown, an arbitrary short pulse width is suggested. After the administration of the initial pulses, the corresponding MEPs *y*_*n*_, *n* = 1, 2, 3 are measured.

In the next stage, an initial estimate of the parameter vector

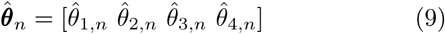

is obtained by fitting the integrated model (1) to the accumulated data set of the baseline and initial pulses for *n* = 3. By using 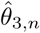, an initial estimate of the membrane time constant, 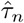 is then computed by solving the following equation for *n* = 3:

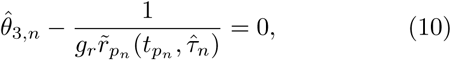

where 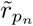 is the normalized depolarization factor after the *n*-th pulse, given 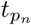 and 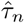, i.e.,

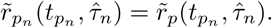

In the next stage, the width and amplitude of the next pulses are chosen in a sequential manner as follows.

The pulse width is tuned by maximizing the depolarization factor per

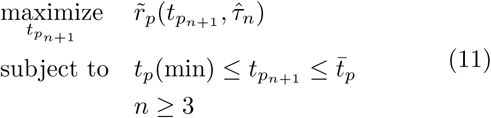

where 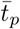 can be set to *t*_*p*_(max) or the critical pulse width 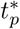 if it is known.

The pulse amplitude is tuned by the FIM optimization as follows:

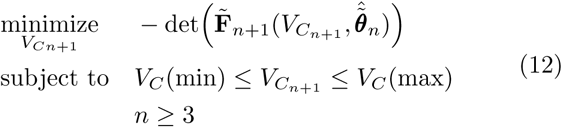

where 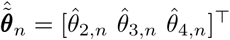, and

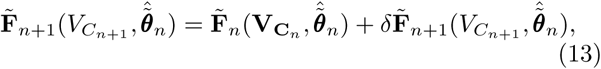

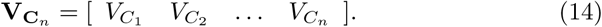

The matrices 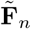 and 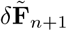 are given in [48].

After the calculation of the next pulse amplitude and width, the pulse is administered and the corresponding MEP is measured. The estimation of the parameter vector and time constant is updated as described above. The process continues until the following convergence constraints are satisfied for *T* consecutive times:

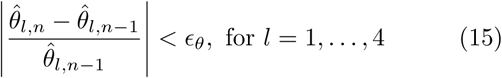

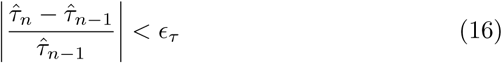

*ϵ*_*θ*_ and *ϵ*_*τ*_ denote the estimation tolerance of ***θ*** and *τ*, respectively. To reduce premature terminations, *T >* 1 is suggested as discussed in [46].

## 5. Simulation Results

Consider 10,000 stimulus–response pairs, shown by ‘×’ in Fig. 4-a, generated by the integrated model (1) with the following arbitrarily chosen true values adjusted to show realistic variability features [55]:

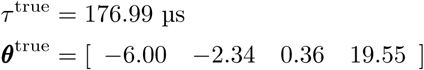

**Figure 4:**
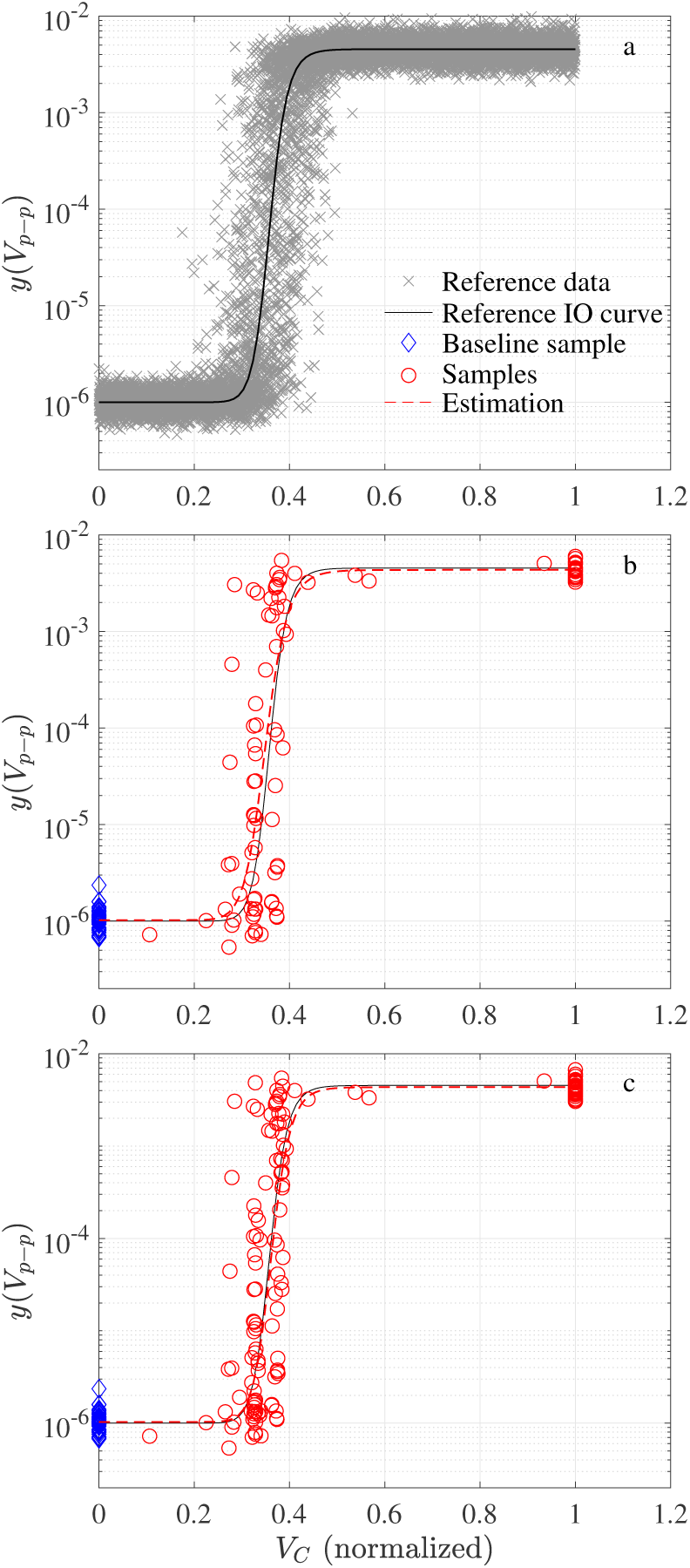
Sample simulation run. (a) reference data and IO curve, and estimation by using the FIM and uniform sampling methods at *n* = *n*_f_ = 99 (b) and at *n* = *n*_max_ = 150 (c).

The problem in this section is to estimate the membrane time constant and reference IO curve by using the proposed automatic tuning of the cTMS pulse amplitude and width and SPE method. From Fig. 2, the critical pulse width is obtained as 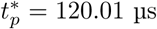. Thus, the integrated model is identifiable for *t*_*p*_ < 120.01 µs.

Fifty baseline samples are arbitrarily taken. The amplitudes of the three initial pulses are arbitrarily chosen 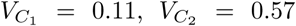, and 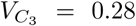, with the same and randomly chosen pulse width at 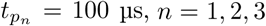, which is less than 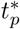 to assure the identifiability. As mentioned, earlier, if 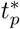 is unknown, a short initial pulse width is suggested.

The trust-region curve fitting algorithm is run with lower and upper limits on the parameter vector as follows:

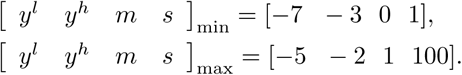

The curve fitting is performed on the logarithmic scale to mitigate the variabilities’ effects [22, 12]. A bad-fit detection and removal technique is employed to improve the curve fitting performance.

The optimization problems (11) and (12), and the non-linear equations (10) are solved by using Matlab fmincon and global search interior-point algorithm with random initial guesses. In (11), the upper bound is chosen as 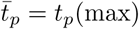 to search the whole range. It could be set to 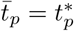 is the critical pulse width is known.

The maximum number of pulses is arbitrarily set to *n*_max_ = 150. The SPE method satisfies the stopping rule at *n* = *n*_*f*_ = 99. Fig. 4-b and Fig. 4-c display the stimulus–repose pairs as well as the estimated IO curve at *n*_*f*_ = 99 and *n*_max_, respectively. As discussed in [46], it is seen that the FIM optimization administers stimuli mainly from three regions, two sectors from the slope area and one sector at around *x*_max_ = 1, which contain the maximum information for curve fitting. Fig. 5 shows the estimation of [*y*^*l*^, *y*^*h*^, *m, s*] with respect to the pulse number. Fig. 6 represents the estimated time constant. The results confirm satisfactory estimation of the reference IO curve and true parameters.

**Figure 5:**
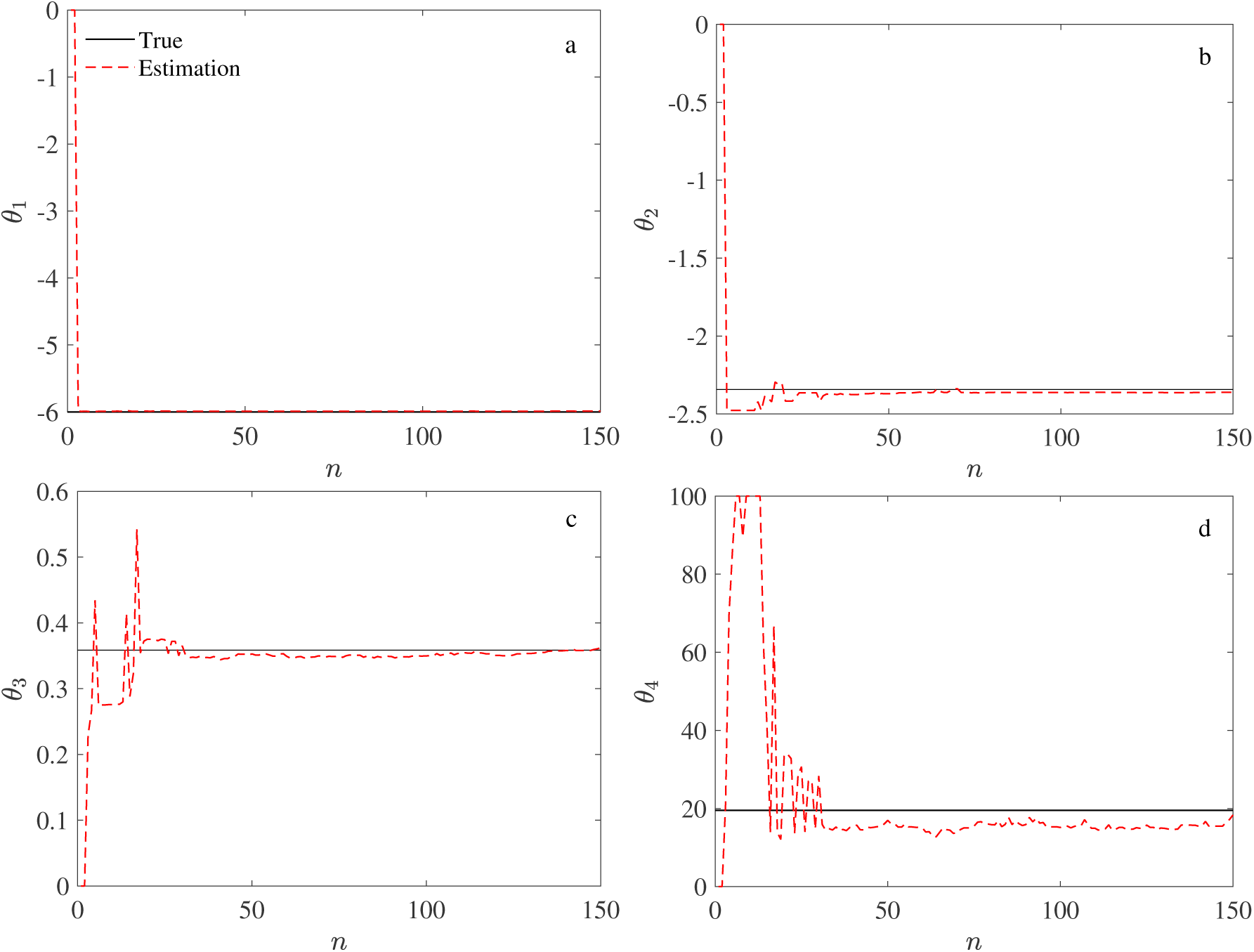
Estimated IO curve parameters. (a) low plateau *y*^*l*^, (b) high plateau *y*^*h*^, (c) midpoint *m*, and (d) slope parameter *s*.

**Figure 6:**
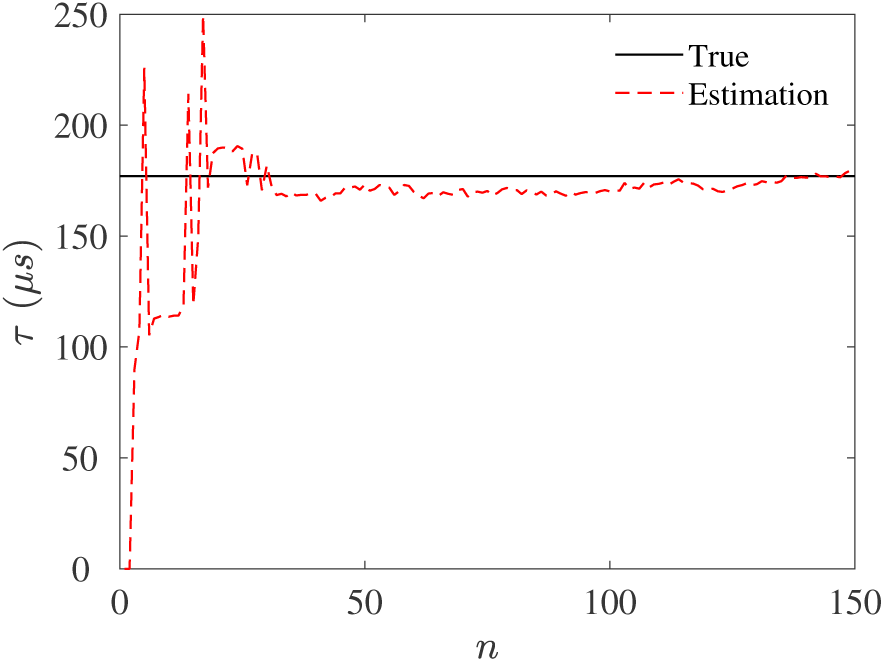
Convergence of the time-constant estimate with increasing number of pulses.

Fig. 7 shows the pulse widths obtained by maximizing the normalized depolarization factor 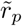. It is seen that 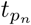 tends to the critical pulse width 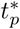. Thus, this method provides a tool for the identification of the critical pulse width 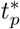 as well, without any prior neurophysiological or anatomical knowledge of the neural membrane.

**Figure 7:**
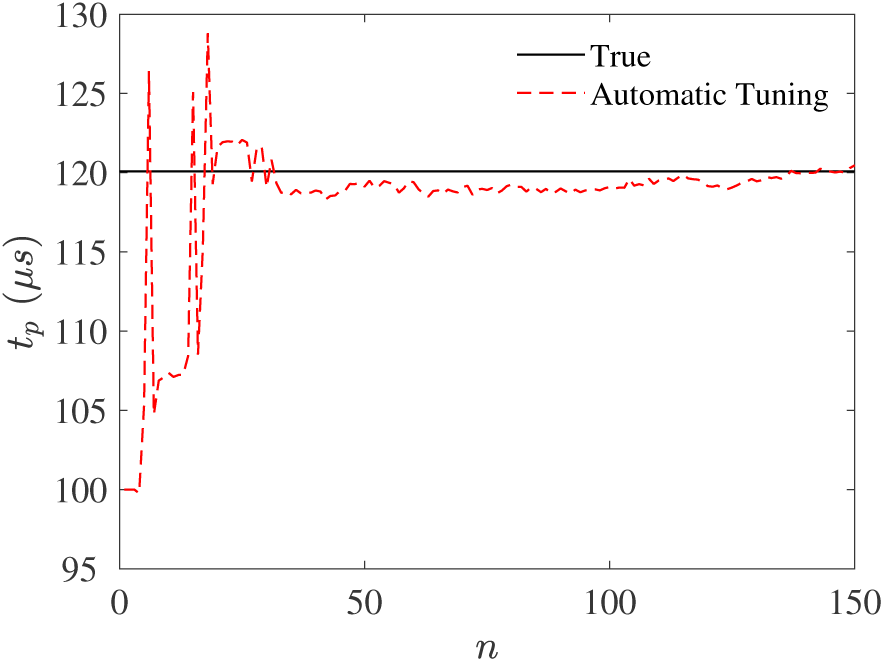
Convergence of the estimate of the critical pulse width with increasing number of pulses.

**Figure 8:**
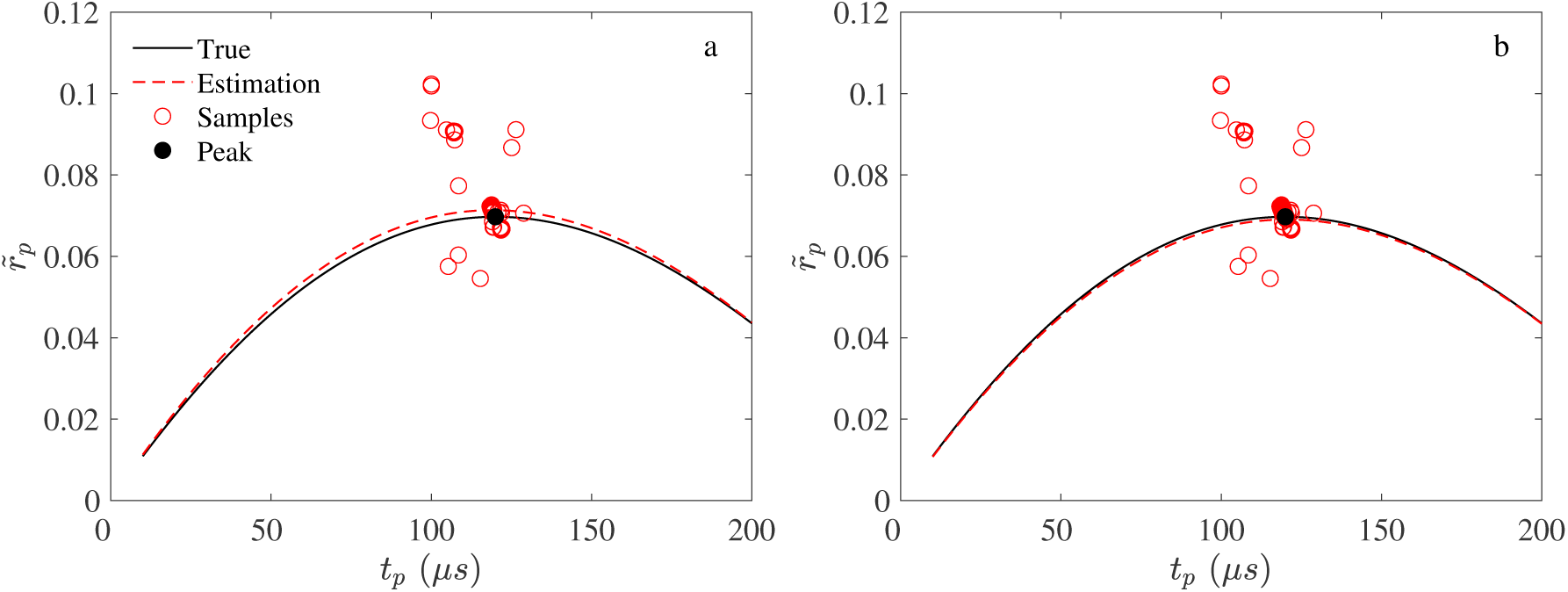
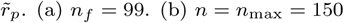. (a) *n*_*f*_ = 99 (b) *n* = *n*_*max*_ = 150.

Finally, Fig. (7) confirms that solving the optimization problem (11) tunes the pulse width to maximize the normalized depolarization factor 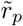.

## 6. Conclusions

This paper confirmed that the optimization of the Fisher information matrix (FIM) and the normalized depolarization factor can serve for automatic and optimal tuning of pulse amplitude and width for sequential parameter estimation (SPE) of the membrane time constant and the input–output curve in closed-loop electromyography-guided (EMG-guided) controllable transcranial magnetic stimulation (cTMS). In addition, the simulation results demonstrated that the normalized depolarization factor maximization can identify the critical pulse width without any prior neurophysiological or anatomical knowledge of the neural membrane.

## Notes

https://github.com/smmalavi/cTMS-Amp-Width-Tuning

### Competing Interest Statement

The authors have declared no competing interest.

